# Honey bee queen production: Canadian costing case study and profitability analysis

**DOI:** 10.1101/2020.01.14.906461

**Authors:** Miriam Bixby, Shelley E. Hoover, Robyn McCallum, Abdullah Ibrahim, Lynae Ovinge, Sawyer Olmstead, Stephen F. Pernal, Amro Zayed, Leonard J. Foster, M. Marta Guarna

## Abstract

The recent decline in honey bee (Hymenoptera: Apidae) colony health worldwide has had a significant impact on the beekeeping industry as well as on pollination-dependent crop sectors in North America and Europe. The pollinator crisis has been attributed to many environmental and anthropological factors including less nutrient rich agricultural monocultures, pesticide exposure, new parasite and pathogen infestations as well as beekeeper management and weather. Canadian beekeepers have indicated that issues with honey bee queens are the most significant factor affecting their colony health. In Canada, beekeepers manage colony losses by relying on the importation of foreign bees, particularly queens from warmer climates, to lead new replacement colonies. Unfortunately, the risks associated with imported queens include the introduction of new and potentially resistant pests and diseases, undesirable genetics including bees with limited adaptations to Canada’s unique climate and bees negatively affected by transportation. Importing a large proportion of our queens each year also creates an unsustainable dependency on foreign bee sources, putting our beekeeping and pollination sectors at an even greater risk in the case of border closures and restrictions. Increasing the domestic supply of queens is one mitigation strategy that could provide Canadian beekeepers, farmers and consumers with a greater level of agricultural stability through locally bred, healthier queens. Our study is the first rigorous analysis of the economic feasibility of Canadian queen production. We present the costs of queen production for three case study operations across Canada over two years as well as the profitability implications. Our results show that for a small to medium sized queen production operation in Canada, producing queen cells and mated queens can be profitable. Using a mated queen market price ranging from $30 to $50, a producer selling mated queens could earn a profit of between $2 and $40 per queen depending on price and the cost structure of his operation. If the producer chose to rear queens for his own operation, the cost savings would also be significant as imported queen prices continue to rise. Our case studies reveal that there is potential for both skilled labour acquisition over time in queen production as well as cost savings from economies of scale. Our queen producers also reduced their production costs by re-using materials year to year. Domestic queen production could be one viable strategy to help address the current pollinator crisis in Canada.

## Introduction

Honey bees play an important role in both natural and managed ecosystems through their pollination services to flowering plants. As such, they contribute substantially to the production of food crops: over a third of global food crop species increase yield as a function of animal pollination, primarily by bees (Klein et al. 2007). In Canada, managed honey bee colonies (*Apis mellifera* L.) contribute to the pollination of many crops including tree fruits, berries, cucurbits, and oil seeds, especially production of hybrid canola seed. In 2016, honey bee contribution to Canadian food crops was estimated at $4-$5.5 billion (HCSDA 2017). Canadian beekeepers managed 803,352 colonies over the 2018-2019 season (CAPA 2019), an increase of over 16,000 colonies from the previous year, however, beekeeper revenues have been decreasing due to falling honey prices (Phipps 2017) and increased colony mortality (CAPA 2019).

Colony mortality has been a concern worldwide for several years, with U.S. beekeepers reporting 38% colony mortality over the 2018-2019 winter, the highest winter loss in recent history (BIP 2019). Canadian honey bee colony winter mortality has also been significant throughout the past decade (Fig 1). Losses of Canadian honey bee colonies over the recent 2018-2019 winter season was 25.7%, ranging by province from 19% to 54% (CAPA 2019). 2018-2019 colony mortality follows the previous year’s losses which reached 32%, the second highest mortality on record since 2008 (CAPA 2019) and more than double the 15% yearly loss that is considered sustainable by apiculturists (Fig. 1) (Furgala and McCutcheon 1992, vanEngelsdorp et al. 2007). Causes of colony mortality are multifaceted (Currie et al. 2010, Potts et al. 2010, vanEngelsdorp 2013) with the predominant factors being queen health and queen age (Genersch et al. 2010, Spleen et al. 2013, vanEnglesdorp 2013, Liu, et al. 2016). In a recent survey, Canadian beekeepers reported that queen issues were the most important factor contributing to colony mortality (Fig. 2). Despite the significant colony losses, beekeepers are able to mitigate high colony mortality by splitting their colonies each spring and installing new queens. These new queens can be reared by the beekeepers themselves, by other local beekeepers or can be imported. Beekeepers can import queens alone, or as *package bees,* which are comprised of 1-1.5 kg of worker bees with a newly-mated queen.

**Figure 1.**
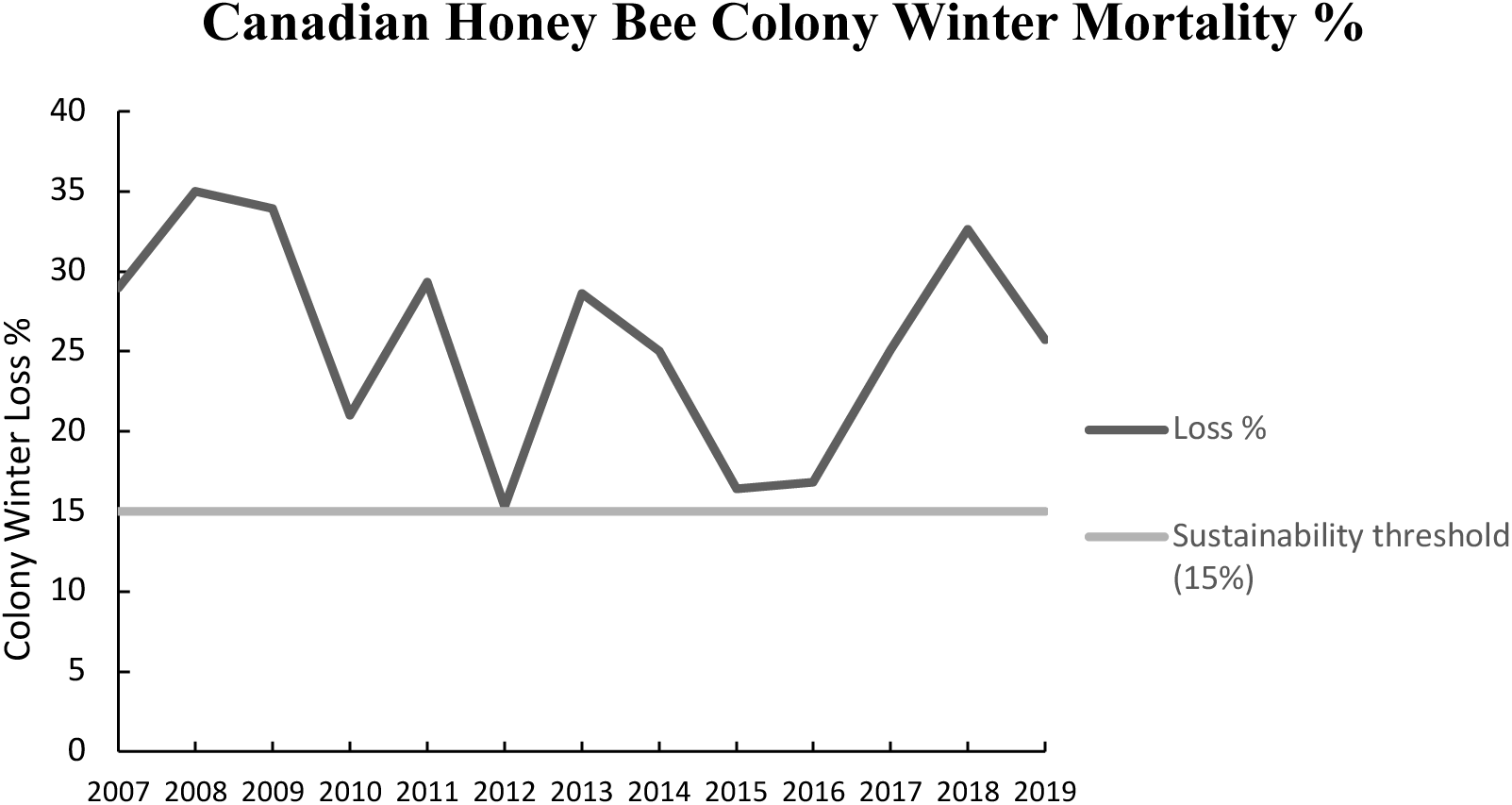
Canadian honey bee colony winter losses over the past decade as reported by the Canadian Association of Professional Apiculturists (CAPA 2019).

**Figure 2.**
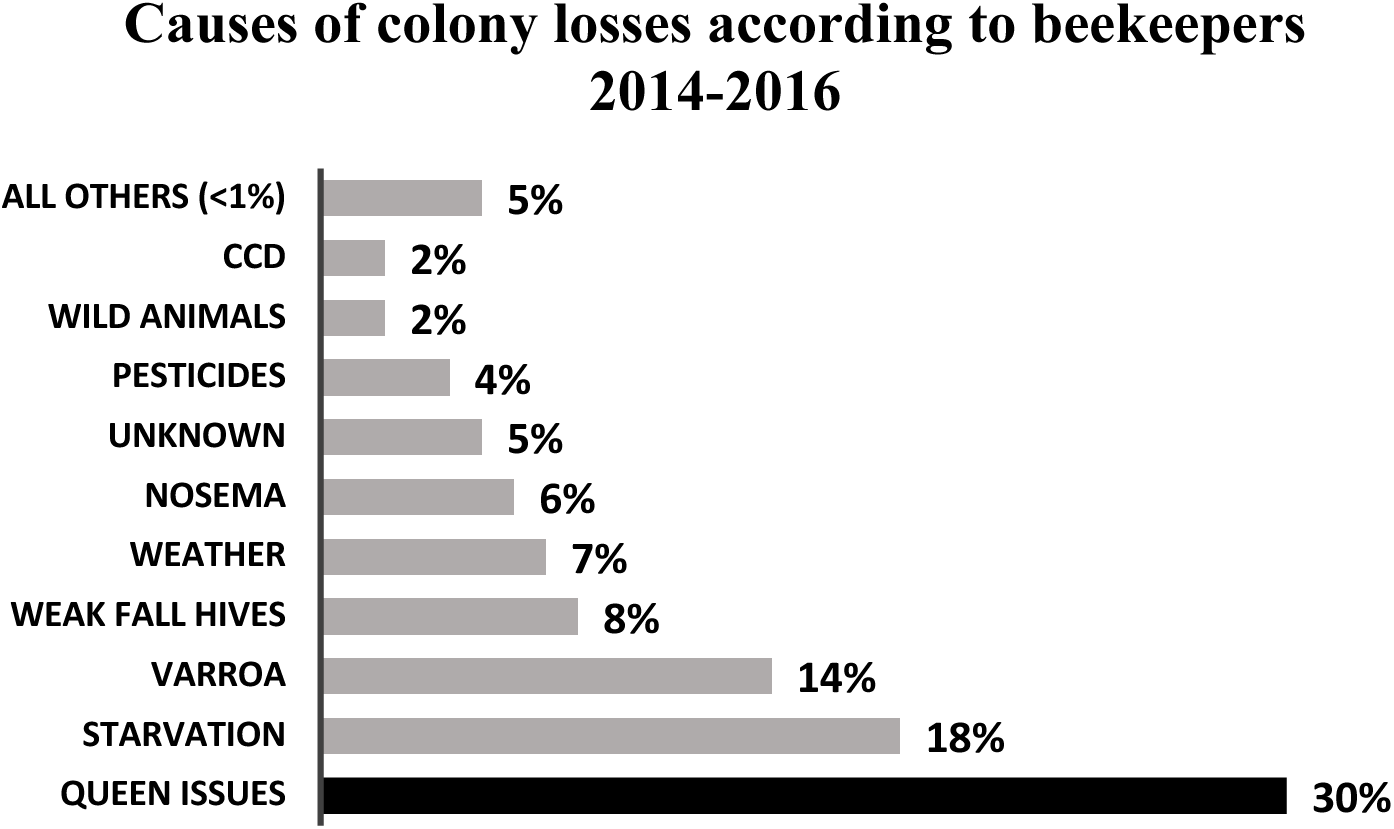
Canadian survey data on beekeeper reported causes of colony mortality in their apiaries through the 2014-2016 seasons (Bixby et al. 2019).

There are an estimated 250-500 beekeepers in Canada who produce queens to supply their own operations and/or sell to other Canadian beekeepers (Bixby et al. 2019). Provincial survey data from 2017-2018 suggests that approximately 100,000 queens were produced in Canada (BCBPS 2016, QIS 2018), a fraction of what is required to support the national population of over 800,000 Canadian colonies (CAPA 2019). Despite the critical role that queen bees play in sustaining Canada’s beekeeping and agricultural sectors, there has been no formal investigation into the economic details of queen bee breeding operations in Canada and no systematic national record keeping of the number of breeders or queens being produced and sold in Canada. Based on survey data (Bixby et al. 2019) and importation statistics (Page, 2017), we know that domestic Canadian queen supply has historically not met demand, particularly in the early spring, and as a result Canada’s beekeeping community has developed a strong culture of queen importation. Large numbers of queens are imported in the spring from warmer climates such as California where breeding can be done much earlier than northern climates. Queens are also imported from regions with contra-seasonal weather such as New Zealand and Australia as well as from aseasonal climates such as Hawaii where queens are reared year-round. In 2018, Canadian beekeepers imported 262,118 queens from Hawaii, California, Chile, Australia and New Zealand (Page 2017) to establish new colonies or to re-queen existing units.

Queen importation, however, is a double-edged sword, simultaneously supplying essential resources for our beekeeping and pollination sectors while risking the introduction of new and potentially resistant pests and diseases, undesirable genetics including bees with limited adaptations to Canada’s unique climate and conditions and/or bees negatively affected by transportation. During transportation, queens can be exposed to temperature extremes that may affect their stored sperm, which in turn can reduce laying success and ultimately impact colony productivity (CFIA 2013, Pettis et al. 2016). Canada’s dependency on foreign queen sources also imposes another potential risk on our beekeeping and other agricultural sectors as prohibitions to importation could result in Canadian beekeepers facing the sudden loss of a quarter of a million queens that the industry is currently unprepared to supply domestically. This is a scenario that Canada narrowly escaped from in 2008 after *varroa* was discovered in Hawaii and again in 2010 after the small hive beetle (*Aethina tumida* Murray), was also found in Hawaii (CAPA 2008, CAPA 2010). Accompanying the risks of importation and the increasing awareness of these risks within the Canadian beekeeping community, has been an unprecedented rise in the prices of imported queen bees from $7.50 in 1988 to $32.50 in 2017, an increase of 333%. Inflation alone accounts for an increase of only 80% (BOC 2019), resulting in a significant real price jump for the beekeeping industry (Figure 3). Adjusting for inflation, real prices rose from just over $12 per imported queen in 1988 up to over $32 per imported queen in 2017. Colony health issues related to imported queens, risks associated with importation, and rising imported queen prices are factors that are concurrently driving an increase in the demand for local queens (Bixby et al. 2019).

**Figure 3.**
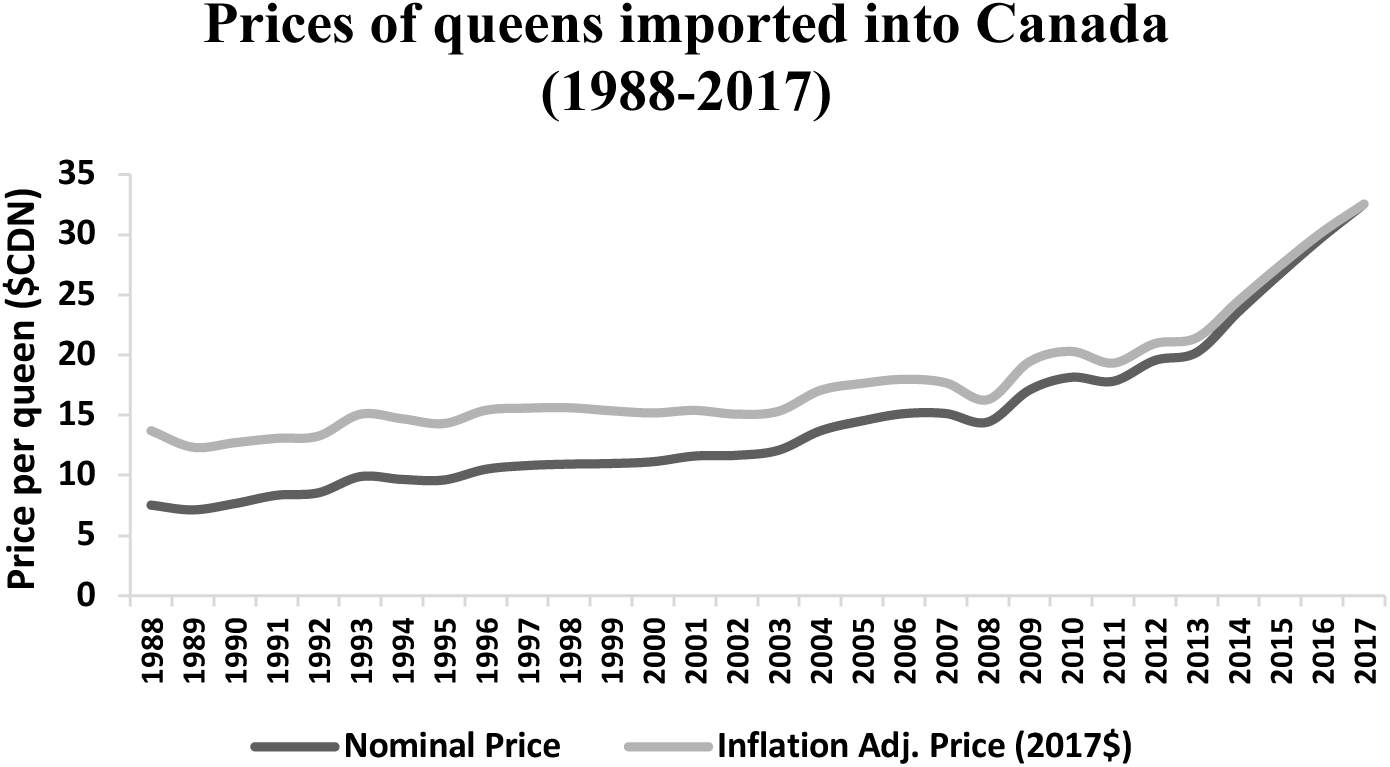
Nominal and real (inflation adjusted) prices for imported queens into Canada 1988-2017 (Page 2017).

Honey bees are social insects with a complex division of labour, which includes the queen who is the sole reproductive female in the colony. The queen mates with between 8-25 drones (males), with an average of approximately 14 drones, over several mating flights (Simone-Finstrom and Tarpy 2018). These mating flight(s) occur very early in her adult life and she stores sperm in her spermatheca for the remainder of her life. To maintain the required worker population, a queen will lay up to 1500 fertilized eggs per day (Winston 1987, Moore et al. 2019), and the resulting female worker bees in the colony are tasked with all non-reproductive colony duties, including caring for the queen, nursing brood, cleaning, and foraging for food. As a result of this matriarchal familial system, the quality of the queen has a direct impact on the colony’s health, productivity and ultimately survival (Nelson and Smirl 1977, Tarpy et al. 2000, Tarpy et al. 2012, Rangel et al. 2013, Simeunovic et al. 2014, Amiri et al. 2017, Eccles et al. 2017). Rearing a queen can involve a rigorous selection process to ensure the new queen carries desirable attributes. This type of selective queen breeding is a specialized skill performed by a small subset of beekeepers. These breeders select for a set of criteria such as honey production, varroa resistance, wintering performance, hygienic behaviour, and/or temperament. Selection usually takes place in the field through specialized phenotypic testing and/or observations of colony performance, however, new laboratory-based testing tools are beginning to reach the market and may soon significantly impact the queen breeding industry in Canada and worldwide (Guarna et al. 2017). These tools would require only a small sample of a colony’s workers to be tested for markers corresponding to specific traits, a much less resource intensive selection process.

As described in Laidlaw and Page (1997), queen rearing requires that the queen producer follow a generalized breeding procedure. Once the queen and drone mother colonies are selected, a process that can be done by the queen producer or within a separate breeding program which then provides the selected genetics to the queen producing beekeeper who uses a queenless cell starter colony to rear queen cells. One-day-old larvae from the selected mother colony are grafted into queen cups and placed into the cell starter colony for the nurse bees to rear (swarm boxes filled with nurse bees are an alternative to starter colonies used by some Canadian producers). After 24-48 hours, depending on the method, the queen cells are moved into a finishing colony (unless using a combined starter-finisher colony) where they will be reared for eight days until they are ready to be sold as queen cells or introduced into small, queenless colonies (mating nuclei) to be mated. Setting up the mating yard(s) requires a significant labour investment and is a critical component in the queen production process. These steps of queen production result in daughter queens that can be used in the originating operation or sold to other beekeepers (Van Alten et al. 2013). Alternatively, a colony can contribute to the production of mated queens by acting as a drone source colony for mating with virgin queens. For the purpose of this manuscript, a ‘queen breeding or production operation’ refers to an operation that is involved in queen production regardless of the method used to select breeder queens.

The risks and costs associated with queen importation can be mitigated by the development of a strong, domestic queen production industry in Canada. Since an important but limited attempt in 1994 (Gates et al. 1994), there have been no rigorous studies delineating the activities of queen breeding operations, assigning breeding and production costs, and examining the profit implications for the industry. In this paper we present the first comprehensive Canadian queen production costing case study. We tracked three domestic beekeeping operations over two years, and explored the profitability of queen production given current prices and various levels of queen production experience as well as variable queen grafting and mating success rates. This study provides the economic foundation necessary to support the expansion of Canada’s queen production sector, providing a sustainable source of queens for our beekeeping and agricultural industries.

## Materials and Methods

We chose three queen breeding operations in Canada each led by an apicultural researcher (with a range of queen production experience) to ensure systematic data collection. Each operation was managed independently and according to the researcher’s own set of criteria. The first operation, OP1, was located near Moncton, New Brunswick in Atlantic Canada where historically there has not been a large honey bee queen production industry. OP1 is itself a large beekeeping operation in eastern Canada that produces several hundred splits each summer with a focus to pioneer rigorous breeding research in eastern Canada using a relatively large number of colonies. The operation was led by apicultural researchers with in-depth beekeeping knowledge but limited queen breeding experience. OP2 was located in Lethbridge, in southern Alberta in close proximity to many commercial beekeepers, and where honey bee colonies are frequently used for canola pollination. OP2 collaborated with two commercial beekeepers with large operations but virtually no queen breeding experience. OP2 was led by a researcher with many years of beekeeping experience, including experience with queen rearing and selective breeding. While OP2 had diverse queen production experience, the beekeepers leading OP2 had collectively less experience than OP3 in large scale queen production. OP3 was located in Beaverlodge, Alberta on the campus of Beaverlodge Research Farm (BRF), Agriculture and Agri-Food Canada, a federal government research facility. The BRF is located in the Peace Region, the center of Alberta’s prolific honey producing region where honey per colony is typically well above the nation’s average of 55 kgs. (Emunu 2017, Page and Darrach 2016). OP3 was a moderately-sized operation led by an experienced queen breeder.

Table 1 lists relevant attributes of the three breeding operations including size, cell and queen numbers, as well as grafting and mating success rates. Grafting success is calculated by the number of successful queen cells in which larvae were successfully reared compared with the number of cups into which larvae were grafted. Mating success refers to the number of emerged virgin queens that are mated (as determined by the queen producer who observes egg laying in the mating colony) compared with the number of virgin queen cells that were introduced into mating colonies or nuclei. Grafting success can be a function of breeder experience, environmental factors and/or resources devoted to the operation. Mating success is a function of emergence rates, weather, and drone sources among other environmental conditions (eg. predation) as well as inherent genetic qualities. Through the springs and summers of 2018 and 2019, the three breeding operations tracked all inputs into both queen cell and mated queen production including bee feed, materials, and labour. Due to the sequential and additive nature of queen cell into mated queen production, inputs into grafting and rearing cells are also included as inputs into mated queens. Thus, mated queen costs are a function of queen cell costs, in addition to costs specific to rearing and mating queens post cell stage. For the purposes of this queen production study, the costs associated with breeder selection are not included in the production costs. Selection and production are two distinct processes and our focus in this paper is to examine the latter. As well, the opportunity costs incurred by beekeepers who invest their labour and beekeeping resources into queen production at the expense of other beekeeping output is not included in these calculations.

**Table 1.** Breeding cost case study operation demographics

|  | Location | Years of intensive breeding experience | Forage | Surroundings | # Queen cells/# cups grafted (2018, 2019) | Grafting Success Rate (2018, 2019) | # Queens successfully mated/# queen cells (2018, 2019) | Mating Success Rate (2018, 2019) |
| --- | --- | --- | --- | --- | --- | --- | --- | --- |
| **OP1** | Moncton, NB | 3 yrs. | Bramble, goldenrod, clover | Somewhat isolated | 359/450, 202/270 | 80%, 75%<sup>1</sup> | 40/60, 80/116 | 67%, 69% |
| **OP2** | Lethbridge, AB | 10 yrs. | Canola, sweet clover | City, other bee yards, ag areas | 36/90, 675/945 | 40%, 71% | 30/36, 356/430 | 83%, 83% |
| **OP3** | Beaverlodge, AB | 15 yrs. | Canola, alfalfa | Isolated from other yards | 125/140, 50/58 | 90%, 86% | 119/125, 50/50 | 95%, 100% |

<sup>1</sup>OP1 conducted their 2019 queen production over two subsequent rounds that are merged together for this costing analysis, however, it is important to note that the grafting success increased from 59% in round 1 to 91% in round 2, indicating potential for rapid skill acquisition for newer queen producers. Potential reasons for low grafting success for OP1 in Round 1 (as self-reported) were identified as: 1) presence of a laying worker in cell builder #2; 2) presence of queen cells in the upper box of cell builders 3) poor grafting technique and 4) weak cell builders.

For this analysis, we are considering only existing beekeepers as viable players to enter the queen production industry due to the high level of skill and beekeeping experience required for queen production, and thus we assume that these beekeepers will use their current operation’s beekeeping equipment such as land, colonies, and bees to conduct their queen rearing. Additional resources used only for cell and queen production including queen rearing materials and feed will be included in the cost analysis for 2018, whereas only additional materials (cell cups, queen cages, feed) that are typically not re-used will be included for year 2. Tables 2a. and 2b. show the inputs and costs associated with cell and queen rearing respectively for all three operations in both years. Table 3 lists pricing and describes the labour activities associated with the labour activity numbers given in tables 2a and 2b. All labour wages are paid at an average of CDN$20/hour to account for both higher skilled labour, less skilled labour and unpaid family labour (Laate 2017).

**Table 2a.**
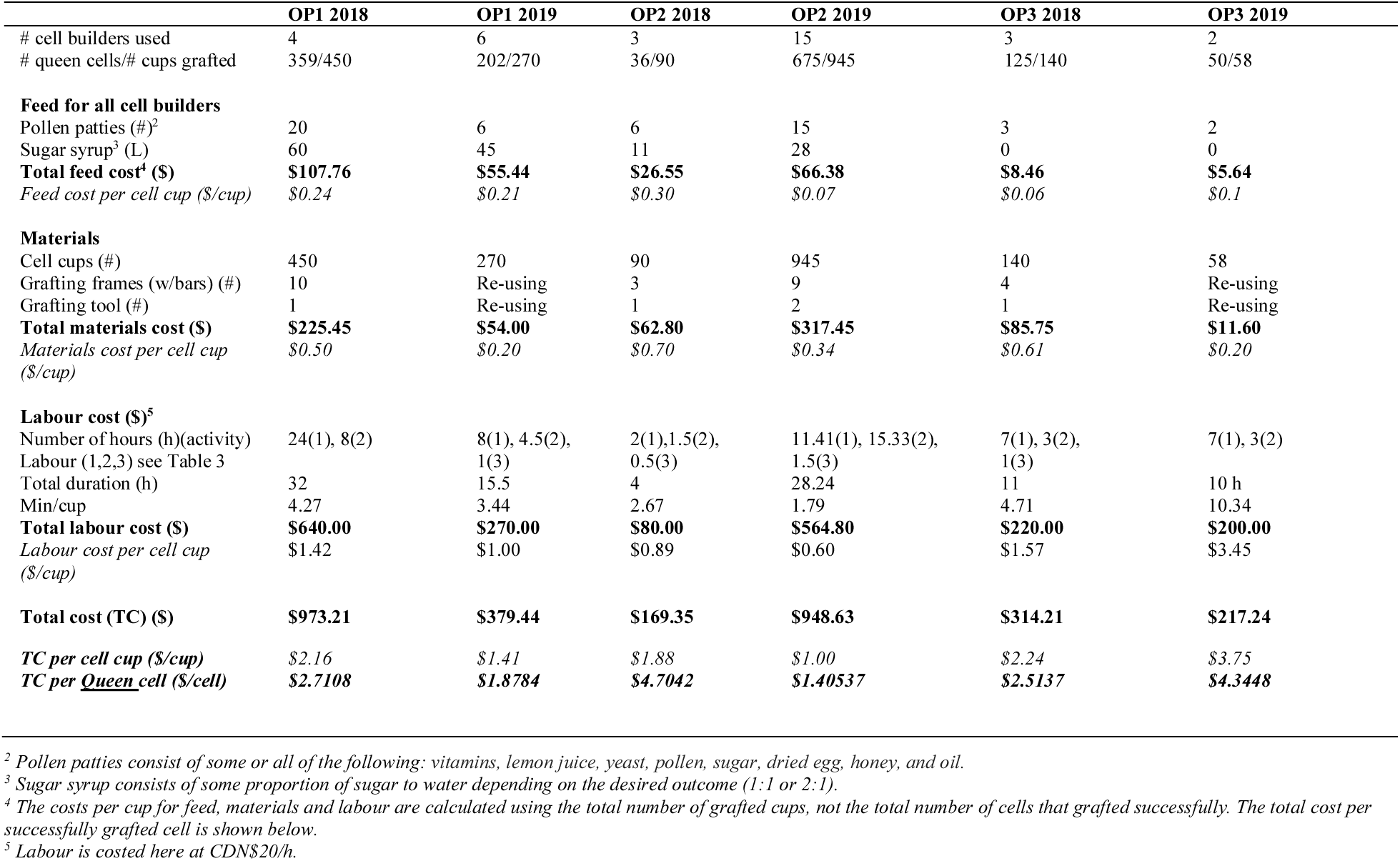
Queen cell breeding costs for three operations

|  | OP1 2018 | OP1 2019 | OP2 2018 | OP2 2019 | OP3 2018 | OP3 2019 |
| --- | --- | --- | --- | --- | --- | --- |
| # cell builders used | 4 | 6 | 3 | 15 | 3 | 2 |
| # queen cells/# cups grafted | 359/450 | 202/270 | 36/90 | 675/945 | 125/140 | 50/58 |
| **Feed for all cell builders** |  |  |  |  |  |  |
| Pollen patties (#)<sup>2</sup> | 20 | 6 | 6 | 15 | 3 | 2 |
| Sugar syrup<sup>3</sup> (L) | 60 | 45 | 11 | 28 | 0 | 0 |
| **Total feed cost<sup>4</sup> (\$)** | **\$107.76** | **\$55.44** | **\$26.55** | **\$66.38** | **\$8.46** | **\$5.64** |
| *Feed cost per cell cup (\$/cup)* | *\$0.24* | *\$0.21* | *\$0.30* | *\$0.07* | *\$0.06* | *\$0.1* |
| **Materials** |  |  |  |  |  |  |
| Cell cups (#) | 450 | 270 | 90 | 945 | 140 | 58 |
| Grafting frames (w/bars) (#) | 10 | Re-using | 3 | 9 | 4 | Re-using |
| Grafting tool (#) | 1 | Re-using | 1 | 2 | 1 | Re-using |
| **Total materials cost (\$)** | **\$225.45** | **\$54.00** | **\$62.80** | **\$317.45** | **\$85.75** | **\$11.60** |
| *Materials cost per cell cup (\$/cup)* | *\$0.50* | *\$0.20* | *\$0.70* | *\$0.34* | *\$0.61* | *\$0.20* |
| **Labour cost (\$)<sup>5</sup>** | | | | | | |
| Number of hours (h)(activity) | 24(1), 8(2) | 8(1), 4.5(2), | 2(1),1.5(2), | 11.41(1), 15.33(2), | 7(1), 3(2), | 7(1), 3(2) |
| Labour (1,2,3) see Table 3 |  | 1(3) | 0.5(3) | 1.5(3) | 1(3) |  |
| Total duration (h) | 32 | 15.5 | 4 | 28.24 | 11 | 10 h |
| Min/cup | 4.27 | 3.44 | 2.67 | 1.79 | 4.71 | 10.34 |
| **Total labour cost (\$)** | **\$640.00** | **\$270.00** | **\$80.00** | **\$564.80** | **\$220.00** | **\$200.00** |
| *Labour cost per cell cup (\$/cup)* | *\$1.42* | *\$1.00* | *\$0.89* | *\$0.60* | *\$1.57* | *\$3.45* |
| **Total cost (TC) (\$)** | **\$973.21** | **\$379.44** | **\$169.35** | **\$948.63** | **\$314.21** | **\$217.24** |
| *TC per cell cup (\$/cup)* | *\$2.16* | *\$1.41* | *\$1.88* | *\$1.00* | *\$2.24* | *\$3.75* |
| *TC per <u>Queen</u> cell (\$/cell)* | *\$2.7108* | *\$1.8784* | *\$4.7042* | *\$1.40537* | *\$2.5137* | *\$4.3448* |

<sup>2</sup> Pollen patties consist of some or all of the following: vitamins, lemon juice, yeast, pollen, sugar, dried egg, honey, and oil.
<sup>3</sup> Sugar syrup consists of some proportion of sugar to water depending on the desired outcome (1:1 or 2:1).
<sup>4</sup> The costs per cup for feed, materials and labour are calculated using the total number of grafted cups, not the total number of cells that grafted successfully. The total cost per successfully grafted cell is shown below.
<sup>5</sup> Labour is costed here at CDN\$20/h.

**Table 2b.** Mated queen breeding costs for three operations

|  | OP1 2018 | OP1 2019 | OP2 2018 | OP2 2019 | OP3 2018 | OP3 2019 |
| --- | --- | --- | --- | --- | --- | --- |
| # Total queen cells used | 60 | 116 | 36 | 430 | 125 | 50 |
| # Successfully mated queens | 40 | 80 | 30 | 356<sup>6</sup> | 119 | 50 |
| **Total cost for queen cells<sup>7</sup>**<br>(# queen cells used) | \$162.6535<br>(60) | \$217.8962<br>(116) | \$169.3500<br>(36) | \$604.3092<br>(430) | \$314.2125<br>(125) | \$217.2400<br>(50) |
| **Feed** |  |  |  |  |  |  |
| Sugar syrup (L) | 227 | 439 | Honey Flow* | Honey Flow* | Honey Flow* | Honey Flow* |
| **Total feed cost (\$)** | **\$192.60** | **\$372.36** | **\$0.00** | **\$0.00** | **\$0.00** | **\$0.00** |
| Feed cost per cell (\$/cell) | \$3.21 | \$3.21 | \$0.00 | \$0.00 | \$0.00 | \$0.00 |
| **Materials** |  |  |  |  |  |  |
| Queen cage (#) | 40 | 80 | 30 | 356 | 119 | 50 |
| Queen candy (#) | 40 | 80 | 30 | 356 | 119 | 50 |
| Marking pen (#) | 1 | 1 | 1 | 2 | 1 | 1 |
| **Total materials cost<sup>8</sup> (\$)** | **\$27.15** | **\$45.35** | **\$22.60** | **\$179.88** | **\$63.01** | **\$31.70** |
| Materials cost per cell (\$/cell) | \$0.68 | \$0.57 | \$0.75 | \$0.51 | \$0.53 | \$0.63 |
| **Labour** |  |  |  |  |  |  |
| Number of hours (h)(activity) | 18(4), 8(5), 10(6) | 14(4), 15(5&6) | 19.25(4), 2(5), 2.5(6) | 109.1(4),<br>51.1(5), 31(6) | 20(4), 4(5),<br>21.5(6) | 11(4), 1.5(5), 3(6) |
| Labour (4,5,6) see Table 3 |  |  |  |  |  |  |
| **Labour cost (\$)** | **\$720.00** | **\$580.00** | **\$475.00** | **\$3830.00** | **\$910.00** | **\$310.00** |
| Total number of hours (h) | 36 | 29 | 23.75 | 191.2 | 45.5 | 15.5 |
| Min/cell | 36.00 | 15.00 | 39.58 | 26.68 | 21.84 | 18.6 |
| Labour cost per cell (\$/cell) | \$12.00 | \$5.00 | \$13.19 | \$8.91 | \$7.28 | \$6.20 |
| **Total cost for Q rearing (\$)**<br>**(including cell costs)** | **\$1102.40** | **\$1215.61** | **\$666.95** | **\$4614.19** | **\$1287.31** | **\$558.94** |
| **Total cost per successfully mated Queen (\$/Q)** | **\$27.56** | **\$15.20** | **\$22.23** | **\$12.96** | **\$10.82** | **\$11.18** |
| **Total additional cost per successfully mated queen (\$/Q) (subtract cell costs)** | **(\$27.56-\$2.71)**<br>**\$24.85** | **(\$15.20-\$1.88)**<br>**\$13.32** | **(\$22.32-\$4.70)**<br>**\$17.62** | **(\$12.96-\$1.41)**<br>**\$11.55** | **(\$10.82-\$2.51)**<br>**\$8.33** | **(\$11.18-\$4.34)**<br>**\$6.84** |

**Table 3.** Materials and labour pricing and description

| Materials (ea.) | Unit Price (ea.) | Labour | Activities |
| --- | --- | --- | --- |
| Sugar syrup (L) | \$0.85 | Cells: | |
| Pollen patty | \$2.82 | 1 | Preparing cells & transporting colonies |
| Cell cup | \$0.20 | 2 | Grafting cells |
| Grafting frame (with bars) | \$12.95 | 3 | Checking cells |
| Grafting tool | \$5.95 | | |
| Queen cage | \$0.45 | Mated Queens: | |
| Queen candy | \$0.005 | 4 | Preparing and transporting colonies and preparing mating yard |
| Marking pen | \$8.95 | 5 | Installing cells and marking queens<sup>9</sup> |
|  |  | 6 | Checking colonies for laying pattern, staff breaks and clean up |

<sup>6</sup> In some queen production operations beekeepers will perform another round of cell introductions to compensate for any poor laying in their mating colonies, this would increase the mating success rate and reduce per queen costs.
<sup>7</sup> To calculate the per cell cost the cost that was incurred to produce each successful cell was used (ranging in Table 1 from \$1.41-\$4.70).
<sup>8</sup> In non-research based operations the queen producer may not use marking pens. In the case of a queen producer using their queens within the operation and thus not for sale, queen cages and candy may not be necessary.
<sup>9</sup> In non-research-based operations, there may not be any labour attributed to queen marking.

## Results

We observed a relative consistency of cell and queen material costs across operations and across time which highlights a systematic cell and queen production process for beekeepers rearing queens and suggests that we may be able to extrapolate these results to a wider queen production sector. The amount of feed per cell builder was up to the discretion of the queen producer and varied greatly between operations. Feed for the mating colonies varied as well but was more a function of environmental factors such as forage availability. OP1 fed the mating nucleus colonies sugar syrup as there was not a sufficient honey flow to provide sustenance for the colonies, unlike OP2 and OP3 who both had strong honey flows at the time of queen rearing and mating. Material costs per cell and per queen were fairly consistent across the three operations in both years. The same materials were used in all three operations and only small differences arose due to the number of grafting frames used with fixed numbers of bars and space for cups. Depending on the number of cups that the researcher chose to graft, some of the equipment was not utilized to full capacity (each frame has three bars and each bar has space for 15 cups) and thus affected the per cup cost. Each operation also had to spread the cost of the grafting tools and pens over the specific number of cells or queens, resulting again in some cost variability. The operations were able to re-use production equipment such as frames, tools and pens, reducing the costs in 2019. Overall, there were minimal cost differentials among operations in per cell/queen materials, however we observed larger differences in per unit feed and labour costs and the three operations also experienced varying grafting and mating success rates.

Labour costs varied tremendously between operations for both cells and mated queen and were a function of breeder experience, management objectives (research-focused operations spent more time with the bees observing specific traits and behaviour for both selection and educational purposes) and the amount of time the breeder was able to allocate to cell and queen rearing that season. As well, there is an economy of scale that develops as the number of queens produced increases while other costs remain static such as travel time to apiaries and some of the general labour involved. These inputs (and associated costs) are incurred regardless of the number of queens, thus as the number of queens produced rises, the per cell or per queen costs decrease. In 2018, OP3 had slightly higher per cell labour costs than OP1, however OP2 had much lower labour costs, a result of the researcher/producer not having much time to allocate to that component of the study. For mated queens in 2018, OP2 had the highest per unit labour costs followed closely by OP1, whereas OP3 had much lower costs, likely a function of streamlining tasks with highly experienced and skilled labour. In 2018, the three breeding operations had a range of overall costs for producing queen cells from $2.51/cell to $4.70/cell and $8.31 to $24.85 for producing a mated queen (Table 4). Total per cell costs for rearing a successful queen cell in 2018 were similar between OP1 and OP3, however, OP2’s overall costs per cell were nearly twice as high as the other two operations, a result of poor grafting success rates which meant higher per cell costs.

**Table 4.** Per cell and per mated queen production costs over three operations during the spring/summer 2018.

|  | Queen Cells |  |  |  | Mated Queens |  |  |
| --- | --- | --- | --- | --- | --- | --- | --- |
|  | OP1 | OP2 | OP3 |  | OP1 | OP2 | OP3 |
| **Feed Cost (\$/cup)** | \$0.24 | \$0.30 | \$0.06 | **Feed Cost (\$/Q)** | \$3.21 | \$0.00 | \$0.00 |
| **Materials Cost (\$/cup)** | \$0.50 | \$0.70 | \$0.61 | **Materials Cost (\$/Q)** | \$0.68 | \$0.75 | \$0.53 |
| **Labour Cost (\$/cup)** | \$1.42 | \$0.89 | \$1.57 | **Labour Cost (\$/Q)** | \$12.00 | \$13.19 | \$7.28 |
| **Total cost (\$/Q cell)** | **\$2.71** | **\$4.70** | **\$2.51** | **Total Additional Cost (\$/mated Q)** | **\$24.85** | **\$17.53** | **\$8.31** |
| | | | | *(minus cell costs)* | *(\$27.56-\$2.71)* | *(\$22.23-\$4.70)* | *(\$10.82-\$2.51)* |

In 2019, the three breeding operations had a range of overall costs for producing queen cells from $1.18/cell to $4.34/cell and $6.84/mated queen to $13.32/mated queen in addition to the queen cell costs (Table 5). OP1 and OP3 reduced the number of cells and queens reared whereas OP2 increased their production of cells and queens between years. The input costs for feed within the operations remained fairly consistent between years which is expected given the management paradigms and availability of forage. However, for OP1 and OP2 there were reductions in materials and labour costs within operations from year to year suggesting both significant efficiency from materials re-use as well as a skill and knowledge acquisition leading to increased labour efficiencies. OP3 experienced an uncharacteristically wet and cold summer with significantly more rain and colder temperatures in 2019 compared to both 2018 and 2017 (GCMCS 2019) making queen rearing more difficult and more than doubling the cost of labour required per grafted cup. As a result, the overall cost to rear queen cells for OP3 nearly doubled from 2018 to 2019. The researcher/beekeeper managing OP3 has extensive queen rearing expertise and thus it would be less likely for OP3 to experience significant skill acquisition and labour cost savings year to year, as labour efficiencies are likely already optimized. Furthermore, given the extreme environmental conditions in 2019 for OP3, the increase in labour costs were not unexpected and in spite of the poor conditions for queen rearing, the experienced beekeeper managed to attain high levels of grafting success.

**Table 5.**
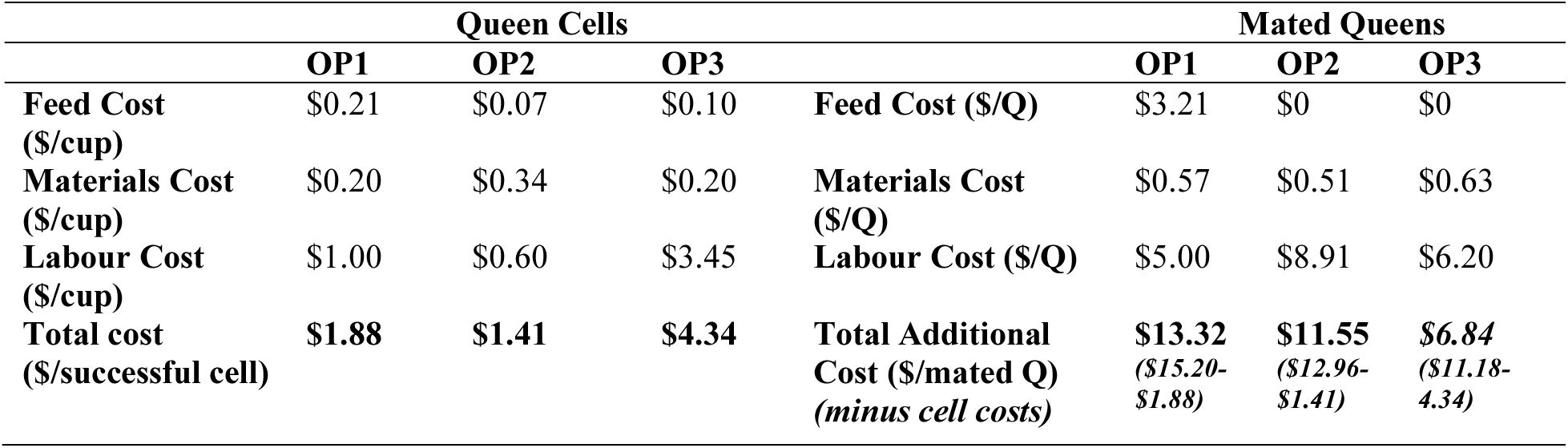
Per cell and per queen rearing costs over two operations during the spring/summer 2019.

Figures 4 and 5 show the overall costs per cells and mated queens as well as the % reduction/increase in costs within an operation between years. Overall cell production costs between two years for OP1 fell from $2.71 per cell to $1.88 per cell, while OP2 saw a reduction in cell costs from $4.70 down to $1.40 over the same two years. As mentioned earlier, OP3 had higher per cell costs in 2019 due to poor weather, however, additional mated queen costs for 2019 for OP3 remained the lowest of the three operations and was even lower than their own additional queen costs in 2018. There were cost reductions for mated queen production between years for all three operations (Fig 5).

**Figure 4.**
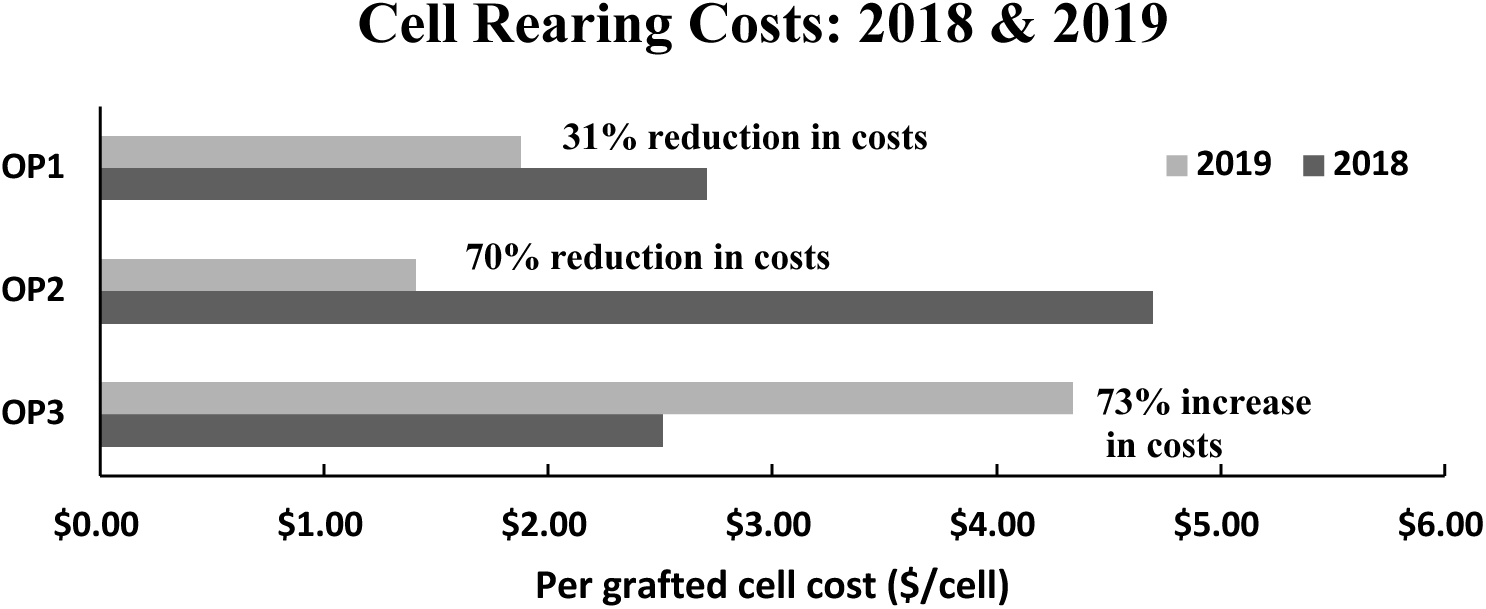
Per queen cell cost differential between first two production years.

**Figure 5.**
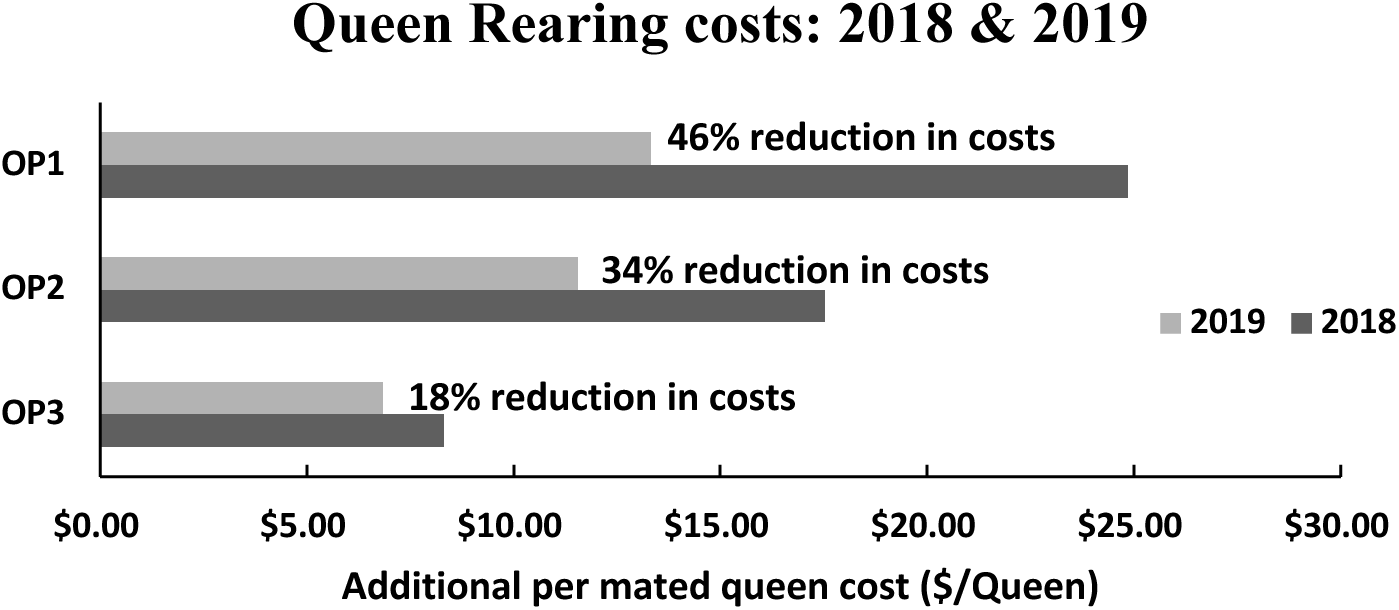
Per mated queen cost differential between first two production years.

As the queen industry in Canada continues to develop and queen producers gain experience and are able to bring costs down, we are seeing alternative queen rearing practices introduced into operations to improve queen and colony health and reduce costs. Some queen producers in Alberta and across the country have begun to introduce queen cells into queenright colonies (colonies with an existing often older and/or less productive queen). This strategy allows the colony to requeen itself as an alternative to producing or purchasing a mated queen, with the same goal of ultimately building-up a stronger, healthier colony led by a young, healthy queen. Mixed success with requeening was reported in earlier research studying the success of introducing queen cells in queenright colonies (Szabo 1982, Jay 1981). However, there is a known positive impact from requeening in terms of decreased winter mortality and increased colony strength particularly when requeening with younger queens (Woyke 1984, Ricigliano et al. 2018), further reducing colony management and replacement costs. It is important that the queen production industry exercise caution, however, in proceeding with this requeening strategy as there is no data to support conclusive positive outcomes (Szabo 1982). The timing for requeening is also of critical importance as there would be a gap in the brood cycle of these colonies. This could negatively impact colony size at critical time points such as pollination contracts, honey flows, or population build up going into winter, unless the requeening method prevents the interruption of egg laying in the colony (Forster, 1972). More research is needed on the biological feasibility and economic efficiency of using cells to requeen queenright colonies.

Introducing queen cells (whether into queenless or queenright colonies) would mean that in the case of our three case study queen producers, an investment of between $2 and $5 per queen cell would potentially yield a strong colony with desirable genetics, saving the queen producer between 77% (OP3) and 90% (OP1) in queen production costs beyond the cell stage (see total additional cost, table 5). In the case of a beekeeper purchasing cells to re-queen colonies, the savings would also be significant as import queen prices continue to rise (Page 2017). In a theoretical example, a commercial beekeeper with costs similar to OP1, investing $2.50 per successful queen cell with a 5000-colony apiary would be able to re-queen half of his colonies for a total cost of $12,500 compared to spending $54,050 (an additional $41,550 beyond the cell stage) to produce 5000 mated queens or purchasing 5000 queens for a minimum of $200,000 ($40 per queen). Alternatively, rather than using their cells or queen in their own operations, queen producers can sell their queen cells and/or mated queens to other beekeepers. In cases of higher overall per cell costs such as in OP2 for 2018, rearing and selling queen cells is less profitable than for OP1 and OP3 due to lower costs, however, for all three operations there are significant profits from selling queen cells in 2018 and 2019. The price for selling queen cells in Canada ranges from $8 to $15 and in some cases even higher (AR 2019, ZQ 2019) resulting in a range of profitability for the three operations at difference price points in 2018. In 2019, due to a decrease in costs (Figs. 4 & 5), we see increases in profitability for both OP1 and OP2 from selling queen cells (Figure 6) and a decrease in profitability for OP3.

**Figure 6.**
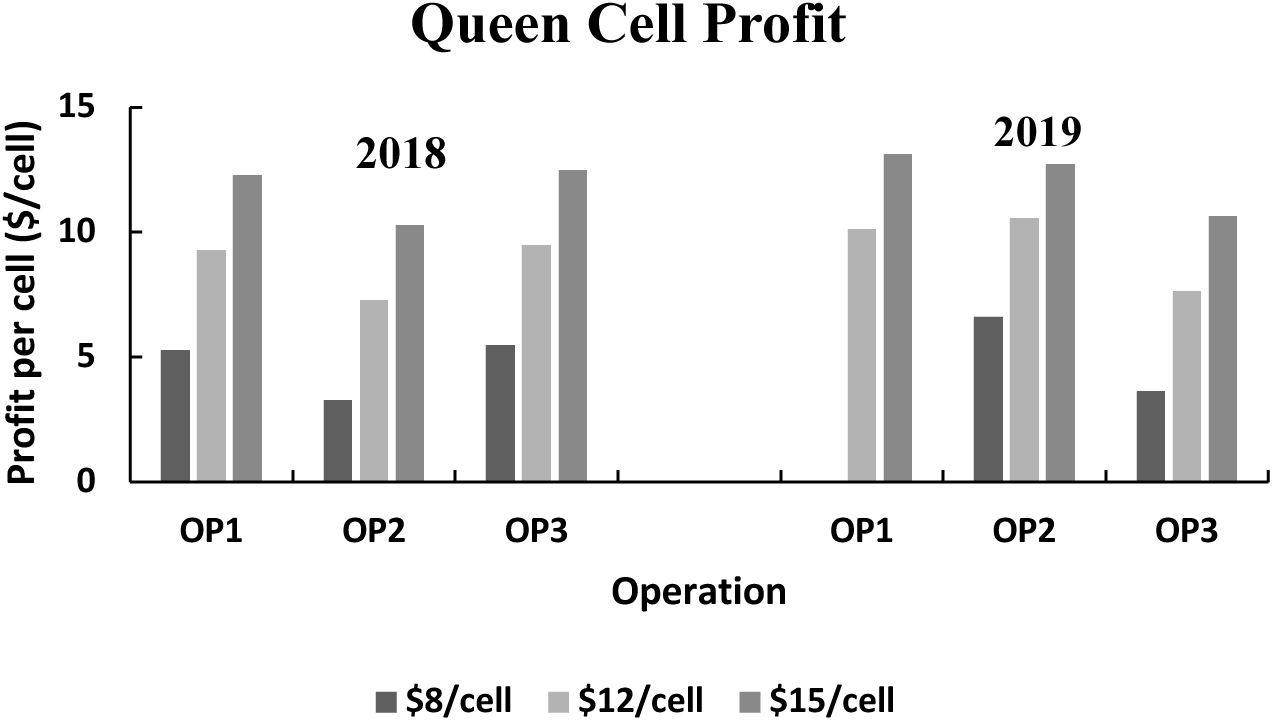
Per cell profit by cell prices and operation costs for 2018 and 2019.

Queen cells require fewer resources and can be produced in larger quantities than mated queens. However, queen cells present a higher risk to the buyer as the queen has not yet emerged or successfully mated, and they are extremely sensitive to transport. Their use increases the risk of queen failure and may require a period of queenlessness for a colony while mating takes place resulting in lost production. As a result of these challenges, demand for queen cells in Canada is much less than the demand for mated queens. Given the recent trend of rising queen prices (Page 2017), the increased demand for local queens, and the willingness of Canadian beekeepers to pay a premium for locally bred queens, domestic queen prices are now in the range of $30-$50/queen (Bixby et al. 2019). These higher prices and the range of mated queen costs for 2018 from $10.84/queen to $27.79/queen, means that all three operations would reap a range of positive profits from just over $2 at $30/queen to over $20 at $50/queen for the high cost operation of OP1. On the lower end of costs, OP3 would reap a profit of $19 for selling a queen at $30 and nearly $40/queen if sold for $50 each. In 2019 we see increases in profit for mated queens for OP1 and OP2. At a price of $40/queen, OP1 sees a 100% increase in profits from 2018 to 2019, OP2 sees a 60% increase in profits while OP3 experiences a small loss of less than $0.40 between the two years (Figure 7). It is important to consider that although these costs take into account queen cages, candy and beekeeper labour, they do not include shipping costs as these can be paid by the receiver or the shipper depending on the contractual agreement.

**Figure 7.**
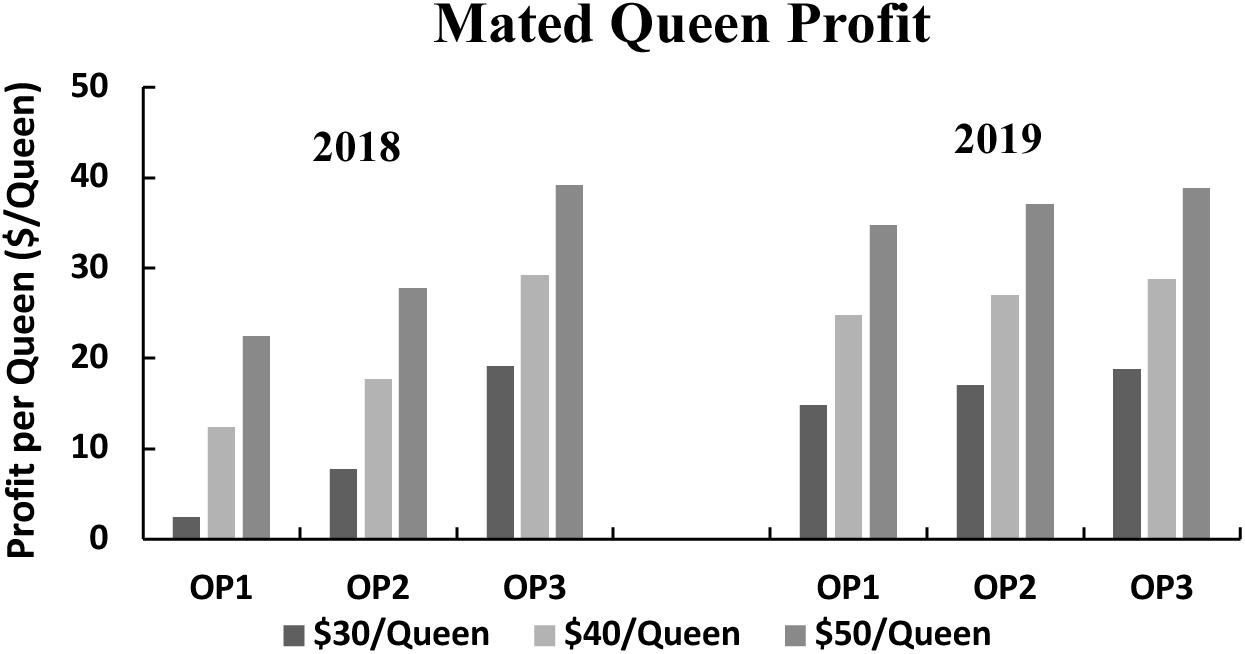
Mated queen profit for three breeding operations given a range of queen prices for 2018 and 2019. Queen costs used for profitability calculations are the full cost of rearing a mated queen including the costs of producing the cells.

For each round of queen production, final per cell and queen costs are highly dependent on both grafting and mating success rates, which varied between operations and over time (Table 1). Figure 8 shows the profitability for mated queens in OP3 in the first, more meteorologically representative year of 2018 given a range of grafting success rates and queen prices, assuming a level of mating success consistent with OP3’s 2018 production year (95%). As grafting success increases with breeder experience and optimal management and environmental conditions (AV 2017, Emsen et. al. 2003), profitability increases although because the impact of grafting success on mated queen costs is relatively small, the increase in profits is also small. For an increase in grafting success from 50% to 75% we see a less than 6% increase in profits and for a jump in grafting success from 75% to 100%, we see an increase of less than 3% in mated queen profits. Mating success has a more significant effect on per mated queen profits. Figure 9 shows the impact that the queen’s mating success has on profitability of mated queens for OP3, given an average grafting success rate for OP3 of 85% and variable mating success rates that are consistent with our three case studies experiences. A rise in mating success from 60% to 80% results in a 19% profitability increase while an increase from 80% to 100% in mating success results in a 10% rise in profits per mated queen. Our researcher-led case studies had variable mating success rates in 2018, ranging from 67% for OP1 up to 95% for OP3 (Table 1), the disparity likely a function of environmental factors. Typical mating success reported in the literature and anecdotally varies between 60-95%. Commercial queen operations typically have high mating success rates as a result of extensive queen rearing experience, skilled labour, and established mating apiaries with proven high success rates.

**Figure 8.**
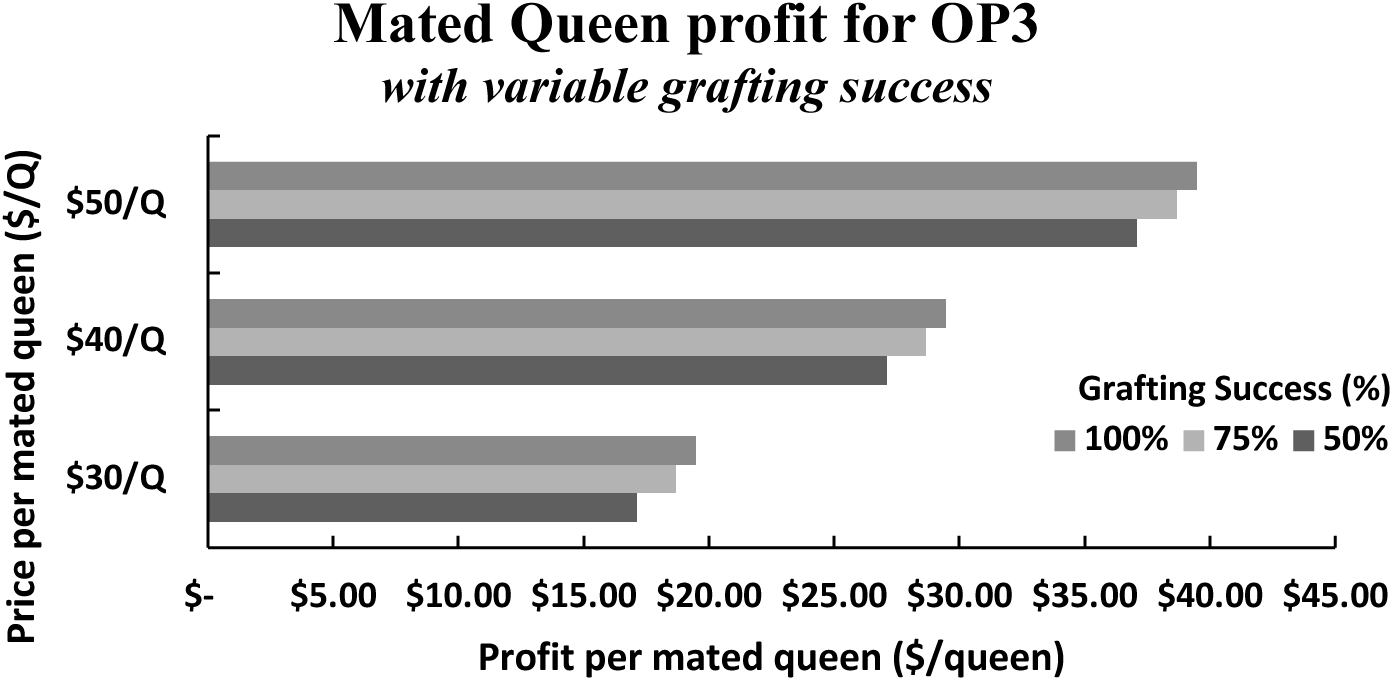
Mated queen profitability given lower production costs of OP3 in 2018, a 95% mating success rate with theoretical grafting success and queen prices.

**Figure 9.**
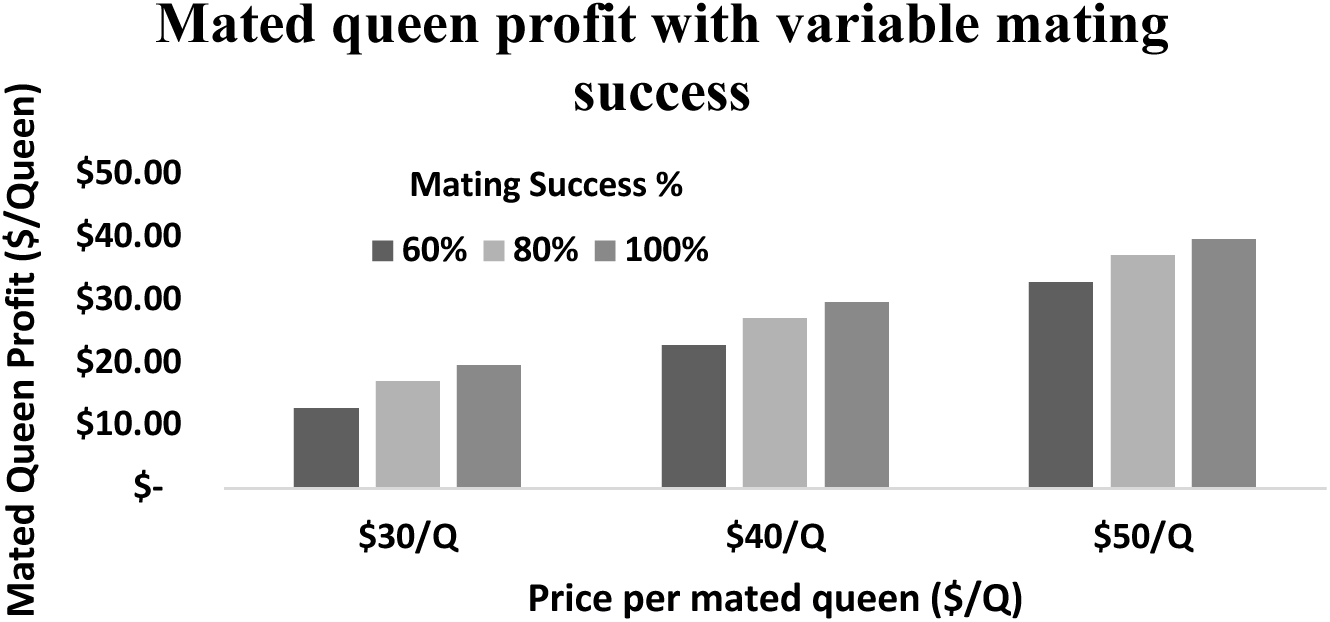
Mated queen profitability given lower production costs of OP3 in 2018, an 85% mating success rate with variable mating success rates and queen prices.

## Discussion

This detailed economic breakdown of Canadian queen production provides evidence that queen production in Canada has the potential to be profitable even for new producers with variable grafting and mating success, as well as when skilled beekeepers are confronted with poor environmental conditions. Based on our study, the difference between total costs and total revenue in the mated queen market in Canada gives queen producers a reasonable profit by absorbing increased costs resulting from any number of factors including environmental conditions and other externalities. For Canadian beekeepers who rear their own cells and queens, there is great potential for cost savings by requeening their own colonies with queens/cells they produce. Whether the beekeeper uses queen cells or mated queens to requeen existing colonies, there will be significant cost savings by reducing queen purchase costs and simultaneously minimizing importation risks to ultimately reduce colony morbidity and mortality.

Each queen production operation in Canada will have a unique approach and expertise with queen rearing which will be reflected in its costs and profitability. However, for queen production, the steps taken and the resources used in each of our three case study operations were determined independently by the breeders without consultation and yet there were similar material costs. From 2018 to 2019, grafting success rates increased along with beekeeper experience. Material re-use and economies of scale for labour were also significant factors in cost reduction and profit increases for operations 1 and 2. Operation 3 had a more uncharacteristic progression from 2018 to 2019 as the climate variability in year 2 posed some significant management challenges and resulted in higher overall cell costs. However, while the experienced queen producer in OP3 likely did not benefit from skill acquisition as such, he did have cost savings from re-using materials in 2019. OP3 was also able to use its queen rearing expertise to mitigate any significant profitability impact from the higher cell costs onto the more salient mated queen production costs. The similarities in material costs between operations reflect a common systematic approach to breeding in Canada, allowing us to extrapolate from our costing analysis to a broader representative Canadian small or medium-scale queen producer and conclude that queen production in Canada has the potential for profit and growth. The three operations’ results in our study offer evidence that small to medium-scale queen production can be profitable. These results likely provide an upper bound for queen production costs, as large-scale commercial queen producers will reap the benefits of even greater economies of scale in their operations, lowering costs even further.

As experienced beekeepers choose to enter the queen production industry, it is important to consider that first year expenditures are higher than in subsequent years. However, even a newly established queen production operation could be profitable given certain environmental and pricing conditions and a skilled beekeeper with some queen experience. Also, as new selective breeding technologies become available to the wider market, Canadian queen production will yield stronger, more highly selected queens that command higher prices. As queen rearing in Canada continues to proliferate and is shown to be profitable, methods will be streamlined further and the number of queen operations and availability of skilled labour should increase, enabling Canadian beekeepers to play a greater role in contributing to this industry’s biological and financial autonomy and sustainability.

## Acknowledgments

Thanks to the following individuals and beekeeping operations for contributing to the queen production in our three case studies: Chris Lockhart with Atlantic Gold/ Lockhart Apiaries and Jillian Shaw (OP1); Jeff Kearns, Rhonda Thygeson, Scandia Honey Co. Super Nuc Apiaries (OP2); Elena Battle, Michael Peirson, Chase Stevens, Jamie Clarke and Carly Balestra (OP3).

This work was part of the BeeOMICS project supported by funding from Genome Canada (227BEE), Genome British Columbia, Genome Alberta, Genome Prairie, Ontario Genomics, Genome Quebec and the Ontario Ministries of Research and Innovation and Agriculture.

## References Cited

Amiri, E., M.K. Strand, O. Rueppell, and D.R. Tarpy. 2017. Queen Quality and the Impact of Honey Bee Diseases on Queen Health: Potential for Interactions between Two Major Threats to Colony Health. Insects, 8(2): 48. doi:10.3390/insects8020048, accessed 18 Nov. 2019.

(AR 2019). Apis Rustica 2019. (http://www.apisrustica.com/rearing.html, accessed 20 Nov. 2019)

(AV 2017). Agriculture Victoria 2017. Raising Queen Honey Bees, 2017. Government of Australia, Victoria State Government. (http://agriculture.vic.gov.au/agriculture/livestock/honey-bees/compliance-and-management/raising-queen-honey-bees, accessed 2nd December 2019)

(BCBPS 2016). BC Beekeeping Production Statistics 2016. BC Beekeeping Production Statistics 2016. British Columbia Ministry of Agriculture, 2016.

(BIP 2019) Bee Informed Partnership 2019. 2018/2019 Total Winter All Colony Loss. (https://research.beeinformed.org/loss-map/, accessed 9 Jan 2020)

Bixby, M., M.M. Guarna, S.E. Hoover, and S.F., Pernal. 2019. Canadian Honey Bee Queen Bee Breeder’s Reference Guide. Canadian Association of Professional Apiculturists Publication, 55 pp.

(BOC 2019) Bank of Canada 2019. Inflation Calculator. (https://www.bankofcanada.ca/rates/related/inflation-calculator/, accessed 6 Jan. 2020)

(CAPA 2008) Canadian Association of Professional Apiculturists AGM Proceedings 2008. Proceedings 2008, Calgary, Alberta. (http://www.capabees.com/shared/2013/02/CAPAProceedings2008.pdf, accessed 7 Jan 2020)

(CAPA 2010) Canadian Association of Professional Apiculturists AGM Proceedings 2010. Proceedings 2010/2011, Markham, Ontario. (http://www.capabees.com/shared/2017/09/2010_11-CAPA-Proceedings-Markham-ON.pdf, accessed 7 Jan, 2020)

(CAPA 2019) Canadian Association of Professional Apiculturists wintering losses. 2019. Annual Colony Loss Reports: CAPA Statement on Honey Bee Losses in Canada: (2007-2019). (http://www.capabees.com/capa-statement-on-honey-bees/, accessed 2nd December 2019)

(CFIA 2013). Canadian Food Inspection Agency. 2013. Risk Assessment on the Importation of Honey Bee (Apis mellifera) Packages from the United States of America (V13) September 2013.

Currie, R. W., F. Pernal, and N. E. Guzman. 2010. Honey bee colony losses in Canada. J. Apic. Res. 49: 104–106.

Eccles, L., M. Kempers, R. M. Gonzalez, D. Thurston and D. Borges. 2017. Canadian best management practices for honey bee health: Industry analysis and harmonization. Bee Health Round Table, Agriculture and Agri-Food, Canada. (http://www.honeycouncil.ca/images2/pdfs/BMP_manual_-_Les_Eccles_Pub_22920_-_FINAL_-_low-res_web_-_English.pdf, accessed 7 Jan 2020)

Emsen, B., A. Dodologlu, and F. Gene. 2003. Effect of Larvae Age and Grafting Method on the Larvae Accepted Rate and Height of Sealed Queen Cell (Apis mellifera L.), J. Appl. Anim. 24:2, 201–206, DOI: 10.1080/09712119.2003.9706457.

Emunu, J. P. 2017. Government of Alberta. Alberta 2017: Beekeeper Survey Results. Alberta Agriculture and Forestry. (https://open.alberta.ca/dataset/a854e8c2-37cf-4c3e-a99f-3bc8e477ca8d/resource/66da7147-0a8c-45ab-8a39-770ce5fd6922/download/alberta-2017-beekeepers-survey-results-final.pdf, accessed 20 Nov. 2019)

Forster, I.W. 1972. Requeening honey bee colonies without dequeening. New. Zeal. J. Agr. Res. 15:2, 413–419, DOI: 10.1080/00288233.1972.10421270.

Furgala B. and D.M. McCutcheon. 1992. Wintering productive colonies. In Graham J M (Ed). The hive and the honey bee (revised edition). Dadant and Sons; Hamilton, IL, USA: 829–868.

Gates, J., J. Howard, and A. Gunner. 1994. Queen Rearing: Spring 1994. Province of British Columbia, Planning for Profit. Agdex 616–810.

(GCMCS 2019) Government of Canada Monthly Climate Summaries 2019 (https://climate.weather.gc.ca/prods_servs/cdn_climate_summary_e.html, accessed 20 Nov. 2019)

Genersch, E., W. von der Ohe, H. Kaatz, A. Schroeder, C. Otten, R. Büchler, S. Berg, W. Ritter, W. Mühlen, and S. Gisder. 2010. The German bee monitoring project: A long term study to understand periodically high winter losses of honey bee colonies. Apidologie 41: 332– 352.

Guarna M.M., S.E. Hoover, E. Huxter, H. Higo, K.M. Moon, D. Domanski, M. Bixby, A.P. Melathopoulos, A. Ibrahim, M. Peirson, S. Desai, D. Micholson, R. White, C.H. Borchers, R.W. Currie, S.F. Pernal and L.J. Foster. 2017. Peptide biomarkers used for the selective breeding of a complex polygenic trait in honey bees. Sci. Rep. 7(1): 8381.

(HCSDA 2017) Horticulture and Cross Sectoral Division Agriculture and Agri-Food Canada 2017. Statistical Overview of the Canadian Honey and Bee Industry and the Economic Contribution of Honey Bee Pollination 2016. (http://www.agr.gc.ca/resources/prod/doc/pdf/honey_2016-eng.pdf, accessed 15th November 2019)

Jay, S.C. 1981. Requeening Queenright Honeybee Colonies with Queen Cells or Virgin Queens. J. Apic. Res. 20(2): 79–83.

Klein, A. M., E. Vaissiere, J. H. Cane, I. Steffan-Dewenter, S. A. Cunningham, C. Kremen, and T. Tscharntke. 2007. Importance of pollinators in changing landscapes for world crops. Proc. R. Soc. Lond. B. 274: 303–313.

Laidlaw, H. and R. Page. 1997. Queen rearing and bee breeding. Wicwas Press; 1st edition. 224 pages.

Laate, E. A. 2017. Economics of beekeeping in Alberta 2016. Economics Section, Economics and Competitiveness Branch, Alberta Agriculture and Forestry. (https://www1.agric.gov.ab.ca/$Department/deptdocs.nsf/all/econ16542/$FILE/Beekeeping2016.pdf, accessed 9 Jan, 2020)

Liu, Z., C. Chen, Q. Niu, W Qi, C. Yuan, S. Su, S. Liu, Y. Zhang, X. Zhang, and T. Ji. 2016. Survey results of honey bee (Apis mellifera) colony losses in China (2010–2013). J. Apic. Res. 55: 29–37.

Moore, Philip A., M. E. Wilson and J. A. Skinner. 2019. Honey Bee Queens: Evaluating the most important colony member. US dept of Agriculture: Cooperative Extension Program. (https://bee-health.extension.org/honey-bee-queens-evaluating-the-most-important-colony-member/, accessed 18 Nov. 2019)

Nelson, D. L., and C. Smirl. 1977. The effect of queen-related problems and swarming on brood and honey production of honey bee colonies in Manitoba. The Manitoba Entomologist 11:45–49.

Page, S. 2017. Statistics Canada. Package and Queen Bee Imports by Source Country by Province, 2017. Canadian Agri-Trade Statistics system (CATSNET).

Page, S., and M. Darrach. 2016. Statistical overview of the Canadian Honey and Bee Industry and the economic contribution of honey bee pollination 2013–2014. Horticulture and Cross Sectoral Division Agriculture and Agri-Food Canada. (http://www.agr.gc.ca/eng/industry-markets-and-trade/canadian-agri-food-sector-intelligence/horticulture/horticulture-sector-reports/statistical-overview-of-the-canadian-honey-and-bee-industry-and-the-economic-contribution-of-honey-bee-pollination-2016/?id=1510864970935, accessed 20 Nov. 2019)

Pettis JS, N. Rice, K. Joselow, D. vanEngelsdorp, and V. Chaimanee 2016. Colony Failure Linked to Low Sperm Viability in Honey Bee (Apis mellifera) Queens and an Exploration of Potential Causative Factors. PLoS One. 2016 Feb 10;11(2):e0147220. doi: 10.1371/journal.pone.0147220. Pettis JS, N. Rice, K. Joselow, D. vanEngelsdorp, and V. Chaimanee 2016. Colony Failure Linked to Low Sperm Viability in Honey Bee (Apis mellifera) Queens and an Exploration of Potential Causative Factors. PLoS One. 2016;11(5):e0155833.

Phipps, R. 2017. International Honey Market Update-June 2017. American Bee Journal. (https://americanbeejournal.com/international-honey-market-2/, accessed 7 Jan 2020)

Potts, S. G., C. Biesmeijer, C. Kremen, P. Neumann, O. Schweiger, and W. E. Kumin. 2010. Global pollinator declines: Trends, impacts and drivers. Trends Ecol. Evol. 25: 345–353.

(QIS 2018). Quebec Institute of Statistics 2018. Main statistics for a few bee products, Quebec. (http://www.stat.gouv.qc.ca/statistiques/agriculture/apiculture-miel/statistiques_principales_produits_apicoles.html, accessed 20 Nov. 2019)

Rangel, J., J.J. Keller, and D.R. Tarpy. 2013. The effects of honey bee (*Apis mellifera* L.) queen reproductive potential on colony growth. Insectes Sociaux. 60(1): 65–73.

Ricigliano, V.A., B.M. Mott, A.S. Floyd, D.C. Copeland, M.J. Carroll and K.E. Anderson. 2018. Honey bees overwintering in a southern climate: longitudinal effects of nutrition and queen age on colony-level molecular physiology and performance. Sci. Rep. 8, 10475. doi:10.1038/s41598-018-28732-z.

Simeunovic, P., J. Stevanovic, D. Cirkovic, S. Radojicic, N. Lakic, L. Stanisic and Z. Stanimirovic. 2014. *Nosema ceranae* and queen age influence the reproduction and productivity of the honey bee colony. J. Apic. Res. 53(5): 545–554.

Simone-Finstrom, M and D. Tarpy. 2018. Honey Bee Queens Do Not Count Mates to Assess their Mating Success. J. Insect Behav. 31(2):200–209.

Spleen, A.M., E.J. Lengerich, K. Rennich, D. Caron, R. Rose, J.S. Pettis, M. Henson, J.T. Wilkes, M. Wilson, J. Stitzinger et al. 2013. A national survey of managed honey bee 2011– 2012 winter colony losses in the United States: Results from the Bee Informed Partnership. J. Apic. Res. 52: 44–53.

Szabo, T. 1982. Requeening honeybee colonies with queen cells. J. Apic. Res. 21(4): 208–211

Tarpy, D. R., J. J, Keller, J.R. Caren, and D.A. Delaney. 2012. Assessing the Mating ‘Health’ of Commercial Honey Bee Queens. J. Econ. Entomol. 105(1): 20–25.

Tarpy, D. R., S. Hatch, S, and D. J. Fletcher. 2000. The influence of queen age and quality during queen replacement in honeybee colonies. Anim. Behav. 59(1): 97–101.

Van Alten, A., J. Tam and R. Bryans., Adapted by: L. Eccles, M. Kempers, D. Rawn, and B. Lacey. 2013. The Ontario Introductory Queen Rearing Manual. Ontario Beekeepers Association, Technology Transfer Program. Guelph, Ontario. Self-published, 70pp.

vanEngelsdorp, D, D. Cox Foster, and M. Frazier. 2007. Fall-dwindle Disease: Investigations into the Causes of Sudden and Alarming Colony Losses Experienced by Beekeepers in the Fall of 2006. Preliminary Report: First Revision, 22 Harrisburg, PA, USA: Pennsylvania Department of Agriculture.

VanEngelsdorp, D., D.R. Tarpy, F.J. Lengerich, and J.S. Pettis. 2013. Idiopathic brood disease syndrome and queen events as precursors of colony mortality in migratory beekeeping operations in the eastern United States. Prev. Vet. Med.108: 225–233

Winston, M. L. 1987. The Biology of the Honey Bee. Harvard University Press. Cambridge, Mass, USA.

Woyke, J. 1984. Correlation and interaction between population, length of worker life and honey production by honey bees in a temperate region. J. Apic. Res. 23:148–156.

(ZQ 2019). Zia Queens Bees 2019. (http://ziaqueenbees.com/queen-cells-nucs-equipment/, accessed 20 Nov. 2019)

